# Carbon ion dosimetry on a fluorescent nuclear track detector using widefield microscopy

**DOI:** 10.1101/2020.03.13.990325

**Authors:** Dietrich W.M. Walsh, Hans Liew, Julian Schlegel, Andrea Mairani, Amir Abdollahi, Martin Niklas

**Affiliations:** Clinical Cooperation Unit Radiation Oncology, German Cancer Research Center (DKFZ), 69120 Heidelberg, Germany; Division of Molecular and Translational Radiation Oncology, National Center for Tumor Diseases (NCT), Heidelberg University Hospital, 69120 Heidelberg, Germany; Heidelberg Institute of Radiation Oncology (HIRO), German Cancer Research Center (DKFZ), 69120 Heidelberg, Germany; German Cancer Consortium (DKTK), 69120 Heidelberg, Germany; Heidelberg Ion-Beam Therapy Center (HIT), 69120 Heidelberg, Germany; Faculty of Physics and Astronomy, Heidelberg University, 69120 Heidelberg, Germany

## Abstract

Fluorescent nuclear track detectors (FNTD) are solid-state dosimeters used in a wide range of dosimetric and biomedical applications in research worldwide. FNTDs are a core but currently underutilized dosimetry tool in the field of radiation biology which are inherently capable of visualizing the tracks of ions used in hadron therapy. The ions that traverse the FNTD deposit their energy according to their linear energy transfer (LET) and transform colour centres to form trackspots around their trajectory. These trackspots have fluorescent properties which can be visualized by fluorescence microscopy enabling a well-defined dosimetric readout with a spatial component indicating the trajectory of individual ions. The current method used to analyse the FNTDs is laser scanning confocal microscopy (LSM). LSM enables a precise localization of track spots in x, y and z however due to the scanning of the laser spot across the sample, requires a long time for large samples. This body of work conclusively shows for the first time that the readout of the trackspots present after 0.5 Gy carbon ion irradiation in the FNTD can be captured with a widefield microscope (WF). The WF readout of the FNTD is a factor ∼10 faster, for an area 2.97 times the size making the method nearly a factor 19 faster in track acquisition than LSM. The dramatic decrease in image acquisition time in WF presents an alternative to LSM in FNTD workflows which are limited by time, such as biomedical sensors which combine FNTDs with live cell imaging.

## Introduction

Fluorescent nuclear track detectors (FNTD) are widely used in radiation dosimetry and radiation biology applications to detect and quantify the energy and trajectory of particles from neutrons to heavy ions [1]. Their usage spectrum spans from quality control of microbeams [2] to neutron dosimetry [3]. One of the most promising current applications in research however is the use of FNTDs for heavy ion beam therapy research. The widespread use of radiation as a therapeutic modality in oncology and the increasing number of hadron therapy centers opening worldwide [4] has lead to the increased interest in the fundamental response of cells to ion irradiation. In the field of fundamental radiation research FNTDs are becoming a truly valuable tool [1]. One such implementation for biomedical radiation research is the **Cell F**luorescent **I**on **T**rack **H**ybrid **D**etector (*Cell-FIT-HD*) biomedical sensor used for in situ biodosimetric detection of single ion traversals in cells irradiated with heavy ions (Fig.1) [5]. The biomedical sensor is based on a Landauer Al_2_O_3_:C,Mg FNTD [6, 7]. Cells are grown on the FNTD and irradiated with ions and their response is then analysed. The ions traversing the cells and the FNTD induce transformation of colour centres visible as track spots along the particles track. These trackspots can be used to reconstruct the particles trajectories through the FNTD and cell layer in order to perform single cell bio dosimetry down to the subcellular level [8, 9]. Cells expressing genetically encoded DNA damage markers such as 53BP1 growing on an FNTD can then be imaged after irradiation to assess the relation of ion hits to DNA foci [10, 11]. Furthermore live cell imaging approaches after ionizing radiation can be used to determine the individual cell fate [12].

The current standard for the dosimetric readout method for the Landauer Al_2_O_3_:C,Mg FNTD is confocal laser scanning microscopy (LSM) [9, 13]. LSM enables precise scanning and detection of fluorescence signal in well-defined optical z-sections. Scanning a high intensity laser over a sample and rejecting out of plane light by an adjustable pinhole before detection by a fast avalanche photodiode is ideal for acquiring signal from the weakly fluorescent tracks in the FNTD. It is however a slow process which limits the size of the field and the resolution to be captured. Due to the high laser intensity and the time required to image a field at a high resolution the LSM is less suitable for experiments where live cells are imaged on the FNTD for days as the cells are more likely to suffer phototoxic damage. In contrast, the ideal imaging setup for live cell imaging of cells growing on the FNTD is a widefield microscope (WF) combined with a high sensitivity sCMOS camera as this enables a larger number of frames to be acquired over time and using a tunable LED excitation. The current readout process of *Cell-FIT-HD* for example uses two microscopes to perform in situ bio dosimetry, the imaging routine used to detect the ionizations in an irradiated FNTD makes use of a LSM and a WF to image the cells growing on the FNTD [14-16].

The readout of the ionization tracks in the *Cell-FIT-HD* experiments is currently performed by LSM after WF imaging of the cells growing on the FNTD has been completed. The imaging of the sample on two microscopes and the consequent image registration of the overlaid field of view (FOV) from WF to LSM adds spatial uncertainty, on the scale of 1 µm [16].

However, the registration of the images between the two microscopes introduces an uncertainty to the ion traversals relative to the cells [16]. The readout of one FNTD field of view stack of 20 images (193×193um) via LSM takes ∼11 minutes. In order to simplify and greatly accelerate the image acquisition and reduce the sources of error the main aim of this work was to use a single WF microscope to enable the imaging of both the cells on the FNTD and the tracks in the FNTD. To ensure the FNTD readout using the WF microscope yielded counts in the expected range for the primary ions, a comparison between LSM and WF, and WF and Monte Carlo simulation were performed. The number of primary carbon ion traversals were imaged and compared with LSM and WF for a near-identical FNTD field. The WF data was then also compared to Monte Carlo (FLUKA) simulations of the irradiation performed at HIT to verify the linear energy transfer (LET) distribution of the carbon ions measured in WF.

## Methods

### Irradiation of the FNTD

Irradiation was performed at the Heidelberger Ionenstrahl-Therapiezentrum (HIT) the setup was the same one as used in [15]. The FNTDs analysed in this paper were irradiated with carbon ions at HIT. The beam energy settings for the accelerator were 122.36-138.66 MeV/u. corresponding to a spread out bragg peak (SOBP) of 1 cm width at 3.5cm depth in water. The total dose delivered to the FNTD was 0.5 Gy. The FNTDs were placed in the centre of the SOBP. The total Monte Carlo (MC) simulated fluence (carbon ions and fragments) was 7.66e+06 ions/cm^2^. The fluence of the carbon ions was 4.16e+06 ions/cm^2^. The dose-averaged LET in water for primary carbon ions was 95.2 keV/µm.

The irradiated FNTD was the physical compartment of the biomedical sensor *Cell-FIT-HD*^*4D*^ introduced in [12]: Briefly, the bottom of a conventional 6 well glass bottom dish (MatTek) was replaced by a disk-shaped FNTD (diameter: 19 mm, thickness: 100µm, Fig.1). The resulting irradiated FNTDs were then stored in the dark until further analysis. The dose uncertainty is taken as 5% for a field of 1×1cm, therefore 0.5 Gy +-0.025 Gy.

**Figure 1.**
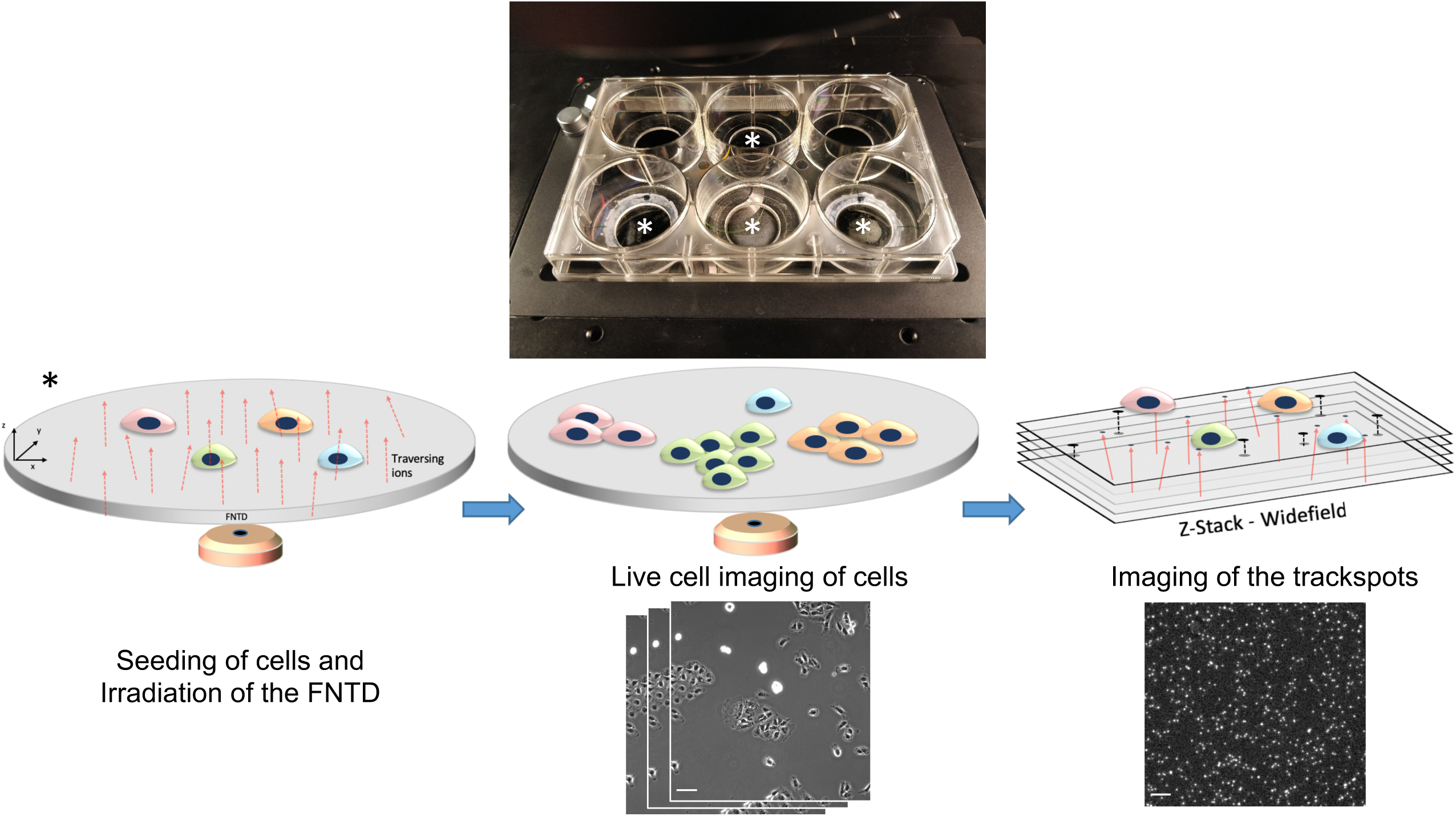
A simplified overview of the biomedical sensor *Cell-FIT-HD*. Upper image is a modified 6 well plate with 19mm diameter FNTDs (FNTD in shape of a wafer 100µm thickness) inserted (*) used for growth of cells during live cell imaging. The three diagrams in the centre and the two corresponding micrographs depict the process of irradiation (left), live cell imaging of cells (centre) and trackspot imaging (including image registration, right). Overview of workflow of *Cell-FIT-HD* and representations of FNTD were adapted from [12]. Scale bars represent 10 µm.

### Microscopy and image analysis

The FNTD readout of the fluorescent trackspots (Ex:620 nm Em:750 nm) was performed on an Olympus IX83 widefield microscope. The IX83 was equipped with a Hamamatsu Orca flash 4.0 v2 sCMOS detector (peak quantum efficiency 82% at 650nm) and a lumencore SpectraX LED illumination system. The sample was excited with a 640/30nm LED at 100% LED intensity (231 mW) using a Plan-Apochromat 40x/1.4 Oil DIC M27 objective. The exposure time was set to 5 seconds and binning 1 x 1, the resolution of the detector was 2048×2048 pixels with an image field of view of 333 x 333 µm. Z-stack height was set to 5 µm. The emission filter was a chroma lumencore X multi-emission filter, the range of detection was 670-760nm. The images acquired were 16bit. Before imaging a flat field correction was performed using a Chroma slide and the Olympus CellSens Software (version 1.15) to ensure flat images.

LSM imaging was performed on a Zeiss CLSM710 with the Zeiss Avalanche Photodiode ConfoCor3 (peak quantum efficiency 63%), Images were captured using a 40x/1.4 Plan-Apochromat oil immersion objective at an image resolution of 1024×1024 pixels with an image field of view of 193 x 193 µm^2^ (zoom of 1.1). The laser intensity was maintained at 100% throughout imaging.

The resulting WF and LSM images were analysed using the FNTD analysis software developed and implemented in ImageJ [13, 17]. The raw images were used in both cases, the previously existing workflow developed for LSM was redesigned so that all of the track analysis parameters apart from threshold were identical for WF. No background correction or signal filtering was performed except for a simple intensity thresholding (MOSAIC Feature point detection parameter: Threshold 0.17 WF and 0.21 LSM) for the individual track spots in the FNTD imageJ macro. The resulting track spots were reconstructed over the z-axis and exported. Histogram Plots of the mean track intensity, the mean value of trackspots along z for each track of valid tracks, were plotted using Rstudio [18].

### Calculation of Trajectory LET

The *LET* of the individual trajectories were determined based on the measured fluorescence intensity of the trajectories, *I*, using the following Equation:

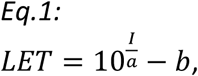

where *a* and *b* are free parameters [9]. In this work, *a* and *b* were optimized manually with the objective to achieve maximum overlap of the measured intensity and simulated LET spectrum in Matlab (R2019b) (Fig.2). The LET values given here are calculated for water. To gain the LET values for Al_2_O_3_ (i.e. in the FNTD) the LET values for water have to be divided by 3.29. Since the form of the LET distribution (Fig. 2) is not altered by this multiplication the LET distribution for water is shown.

**Figure 2.**
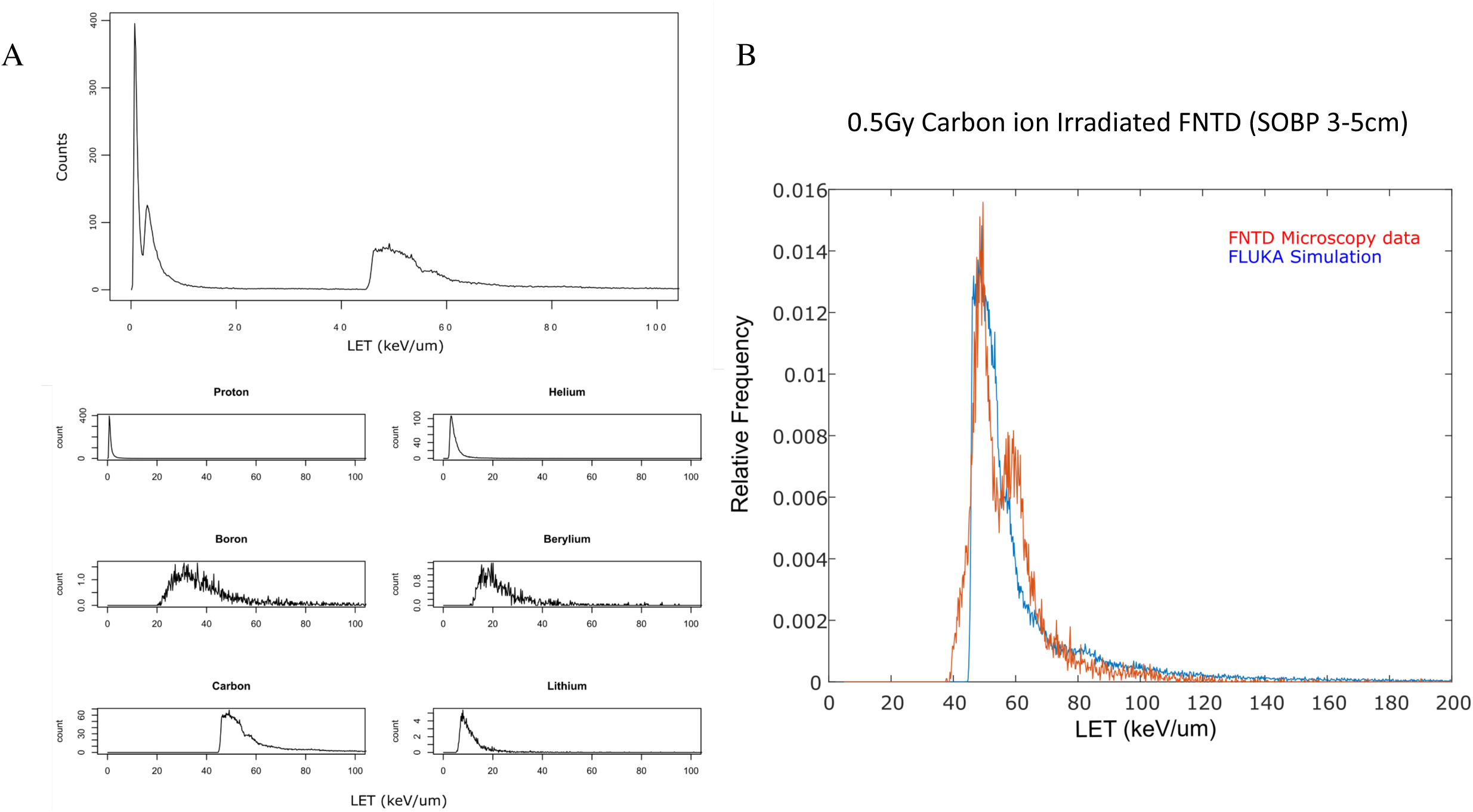
Comparison of Monte Carlo (MC) simulation and read-out by WF. (A) MC simulation (FLUKA) for 0.5 Gy irradiation (C-12, SOBP), including all secondary particles (except electrons). The FLUKA simulation shows the whole LET (Linear energy transfer) spectrum of ions proton, helium, boron, beryllium, carbon and lithium for an area of 333×333 um^2^ (a microscope FOV). The six individual graphs below show the counts expected in the 333×333 µm area per ion type. (B) LET spectra of carbon ions gained by FLUKA (blue) and the recorded data from the microscopy read-out (red) are shown. The two curves fitted using Eq.1 have a high degree of overlap indicating a good correlation between the simulated data and the recorded results.

### Monte Carlo simulation (FLUKA)

Monte Carlo simulation for the Carbon ion irradiation performed at HIT was simulated for a Spread out Bragg Peak (SOBP) at 3.5cm depth and a field of about 13 cm x 9 cm. The FLUKA Monte Carlo version fluka.2011.2x.8 has been used. The default settings employed for hadron therapy application have been used as described in [15, 19]. A detailed geometry of the HIT beam-line was implemented. In order to get enough statistics, about 1% of the planned particle fluence has been simulated, i.e. 7.66e+06 ions/cm^2^. In order to access the information (charge, mass, energy per nucleon and LET) of the particles crossing the detector, we have implemented an event-by-event read-out of MC simulations allowing us to use the same reconstruction constraints in both the simulation and in the experimental data. The Monte Carlo simulation of the particle irradiation performed in FLUKA yielded a total particle fluence of 1.54e+07 for 1 Gy. The fluence consisted of carbon, beryllium, boron, helium and proton counts. For LET calculations in water, for each particle traversal (Z,A) at a certain energy per nucleon, the corresponding LET value is sampled on-line using FLUKA internal routine.

### Direct track comparison between WF and LSM

To allow direct track correlation of the LSM and WF data sets an overlapping FOV was extracted from both datasets. For this purpose the LSM data was rescaled and translated to gain spatial overlap of WF and LSM images. Spherical aberrations at the edges of the images occur in the LSM images, interfering the direct track analysis. The LET data set was thus cropped. 148 pairs of identical ion tracks in the LSM and WF data sets were chosen and compared. The mean intensities of 148 identical tracks detected by the particle tracker in the WF and LSM data sets were compared. The distance between the paired tracks was at maximum three pixels (0.48um WF, 0.56um LSM). To minimize the probability of false positive correlation by intersecting ion tracks, the 148 track pairs were selected manually. For comparison of track intensities in the WF and LSM data sets, the relative difference *ΔI* was calculated by

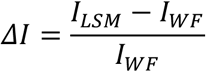

With *I*_*LSM*_ and *I*_*LSM*_ being the track intensities detected by the FNTD analysis software.

## Results

FNTDs Irradiated with 0.5 Gy carbon ions were imaged using the WF microscope and the data analysed to determine the track spots of the primary ions (carbon ions) using the FNTD analysis package [13]. The first image stacks captured with WF showed that it was possible to detect and reconstruct ion tracks using the widefield system (Fig.3). Ionization tracks created by the carbon and secondary ions (Be, B, He, P) in the irradiation field in the FNTD were visualized with the WF system. When compared to FLUKA simulations it was clear that neither the LSM or the WF were capable of detecting all of the secondary particle species which were expected to be seen based on the FLUKA simulation. As the dose deposited in the cells by the primary carbon ions accounts for 93.4% (according to FLUKA simulation) of the total dose it was therefore decided to focus on the detection of the carbon ions as these cause for the majority of the dose and have can readily be detected. MC simulated carbon ion fluence for the 0.5 Gy irradiation plan was calculated to be 4.165e+06 cm^-2^, in comparison the measured fluence in the FNTD as measured by WF was 4.452e+06 cm^-2^ (mean of three 333×333 µm FOV images measured) there is therefore still an overestimation off around 6.8%.

**Figure 3.**
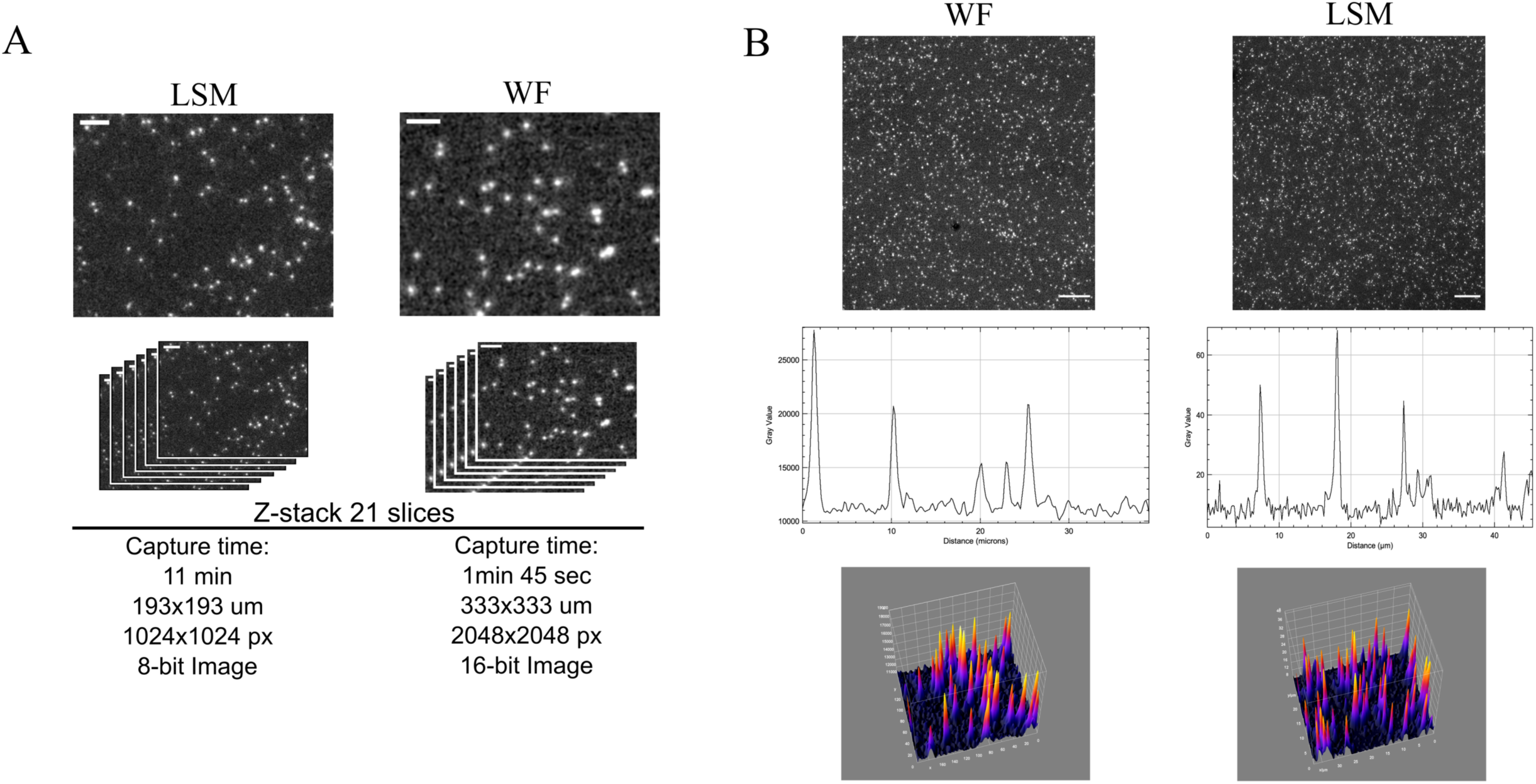
Overview of the image stacks captured using the LSM and WF microscopes of the same 0.5 Gy carbon ion irradiated FNTD. (A) Direct qualitative comparison of image stacks taken with confocal and WF microscopes, two stacks with 21 slices were captured, the details are listed below each image. (B) A representative image captured with LSM and WF using 40X objectives (upper).The line plots show representative signal peaks and background measurements across a cental section of the FOV. The 3D signal peaks are taken from the centre of the FOV and show a similar morphology. The LSM image is 8 bit and the WF image is 16 bit. The scale bars represent 5 and 10 µm respectively in A and B.

The WF datasets were not directly comparable to the images produced by the LSM due to the major differences in microscope systems. LSM works by point scanning whereas WF captures the whole field with a single image. The maximum intensity projection of the recorded track spots in the identical region of the same FNTD imaged with both LSM and WF shows a very high degree of spatial overlap (Fig. 4A). A total of 484 and 571 ion tracks were reconstructed by the FNTD analysis software in the WF and LSM imaging data in this field, respectively (Fig. 4D). Adjusting the corresponding intensity-based threshold for the separation of primary and secondary ions, 425 and 418 carbon ion tracks were identified in the WF and LSM imaging data respectively. The discrepancy of the total number of ion tracks in both data sets arises from the detection of secondary ions. By LSM a factor of 2.6 more secondary ions could be detected. The corresponding intensity distributions of the C-12 tracks are quite similar in both data sets (Fig. 4B). Despite the inherent differences in microscope systems the intensities of 148 identical ion tracks in the WF and LSM data were compared: the median relative difference was 12% (25^th^ and 75^th^ quantile: 10% and 15%, Fig. 4C).

**Figure 4.**
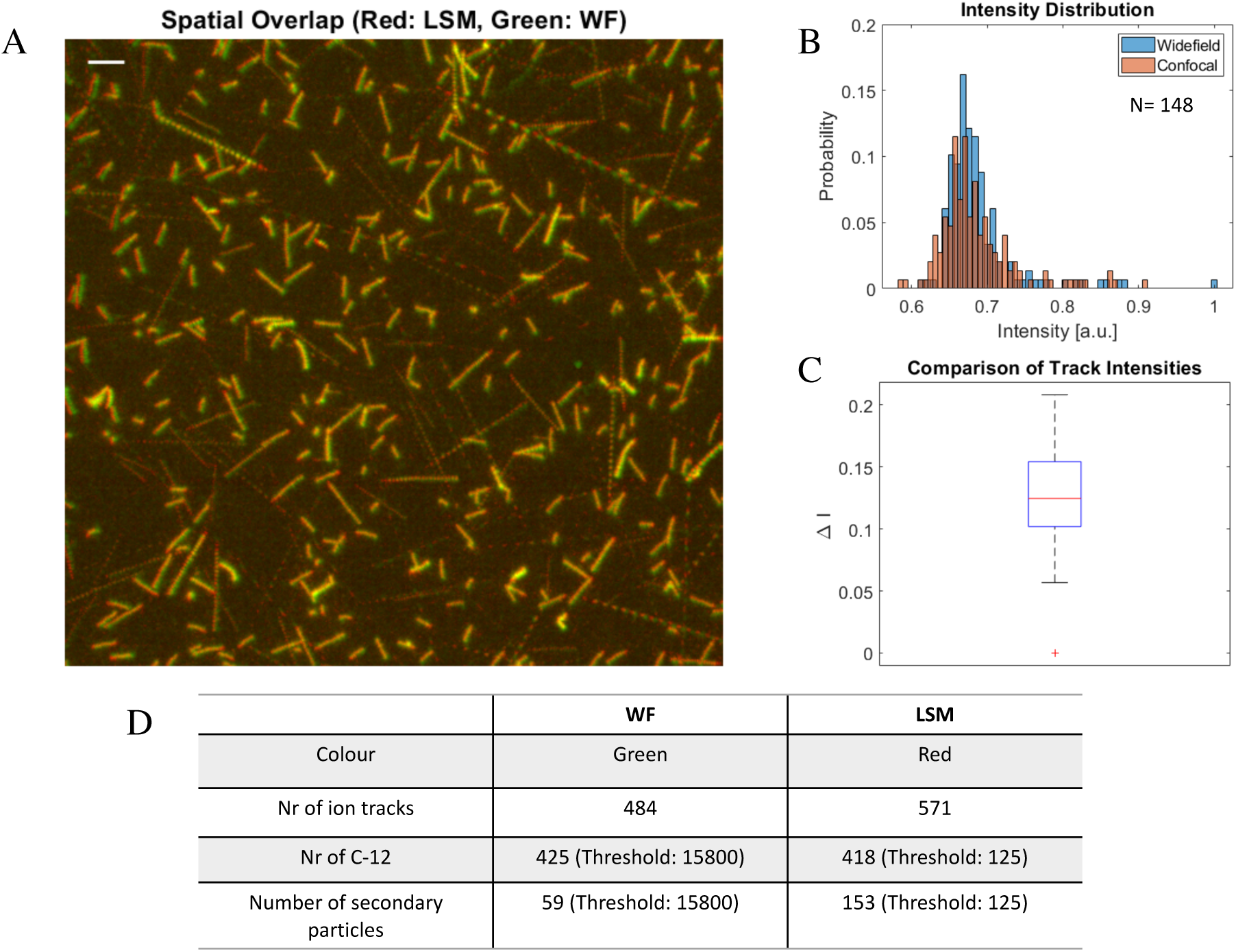
Comparison of tracks detected in the FNTD by LSM and WF. (A) Spatial overlap of read-out signal by WF (green) and LSM (red). The spatial overlap for primary particles is therefore yellow. Maximum intensity-projection is shown. Scale bar, 10µm. (B) The normalized intensity distributions for primary particles gained by WF (blue) and LSM (red) read-out show a high level of overlap. The intensity values were normalized based on the maximum intensity value due to the difference in bit values in each image (8 bit vs 16 bit). (C) The corresponding box and whisker plot shows the relative difference in track spot intensities recorded by LSM and WF (Plot Parameters: Median, 25 and 75 percentile, min max and outlier (red cross)). (D) The table shows the number of ion tracks extracted from the overlap image in (A), with their defining intensity-based thresholds to discriminate between primary ions and fragments.

The exposure time on the WF was increased to 5 seconds to ensure the largest possible dynamic range of the detector could be used while reducing acquisition time. The use of a high quantum efficiency and low dark current sCMOS detector on the WF microscope enabled the more rapid acquisition of the z-stack (21 slices) than possible on the confocal for a larger field of view. The difference in acquisition time was nearly a factor 10 faster, 1 minute 45 seconds for WF and 11 minutes for the confocal stack. The area acquired by WF is in addition 2.97 time larger than that of LSM. Therefore the acquisition of the tracks in the wafer can be acquired up to ∼19x faster using WF.

The main difference in the analysis step is the trackspot threshold definition in the FNTD analysis software, which is set for each imaging method separately. The images taken on the WF and LSM can be analysed by the FNTD package in their raw unaltered state without any correction.

The reconstruction of the ion tracks using the FNTD imageJ package shows that the quality of the WF images were suitable for an analysis of the particle tracks, although the package was developed for LSM images. The data were thresholded during track analysis to determine the detection of primary and secondary particles. The higher the threshold value the less likely that secondary particles are tracked during the analysis. The threshold used was integrated in the FNTD package, the three chosen threshold values were plotted in Rstudio (Fig.5). A threshold value of 0.17 was determined to minimize the secondary particle detection without significantly reducing the number of primary particles detected.

**Figure 5.**
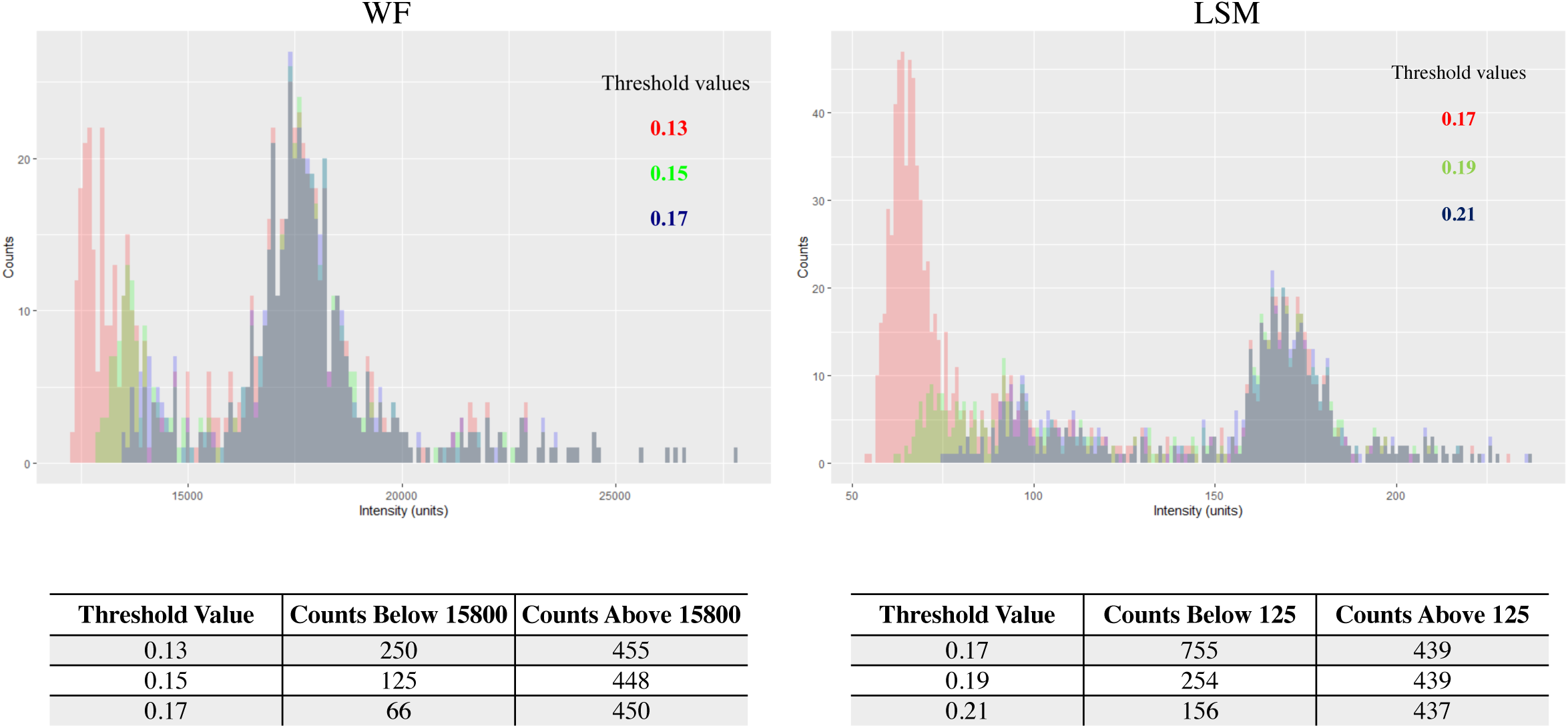
Histograms of three different threshold values used for carbon ion track reconstruction threshold testing on WF (Left) and LSM (right). The graphs show the distribution of track intensity and frequency for varying threshold values (WF; 0.13-0.17, LSM 0.17-0.21 FNTD analysis software). When analyzing the tracks in the FNTD with the FNTD package in imageJ a MOSAIC threshold value must be set, to ensure the least amount of secondary particles are included in the analysis. To define an ideal threshold, different thresholds were tested and the resulting track spot histograms plotted in Rstudio. The graphs show that with increasing threshold value the secondary peak, left in all graphs, shrinks as the value is increased while the primary peak, right, stays unchanged. The value of 0.17 (WF) and 0.21(LSM) were chosen for further Primary particle analysis.

To verify the LET distribution of the primary carbon ions, the extracted LET values (by the FNTD analysis software) from the WF images were compared to the results from FLUKA simulation data. To compare the two distributions the raw FNTD track data were analysed and the threshold was set (value: 0.17) to exclude the majority of secondary ions without changing the distribution of primary ions.

The simulation of the particles depicted in Fig. 2A shows that although the majority of detected particles in the LET range of 45-100 Kev/um are carbon ions there is also a small chance of detection of Boron and Berylium, therefore the measurement of primary ions may also include these ions. The FLUKA simulation data for primary carbon ions was calculated for 1 cm^2^, this was first normalized to the area microscope FOV (333×333) then using Eq.1 the LET of the individual trajectories was calculated and (measured track intensity vs FLUKA) were plotted in Matlab (Fig.2B). The resulting graph shows a high degree of overlap of the two curves and a calculated dose by the FNTD analysis software of 0.52 Gy (carbon ions) from the measured track spots. The applied physical dose for the FNTD was 0.5 Gy (SOBP, all particles) with a dose uncertainty of 5%. The dose contributed by carbon ions alone was calculated to be 0.467 Gy (FLUKA calculation; carbon contributes 93.4 % of the dose). The detection mismatch between the planned physical dose of carbon ions (FLUKA) and the detected carbon ion dose (FNTD) for one analyzed field (333×333 µm) is therefore 6.8% for the settings chosen. In addition to the fast readout time and ability to detection primary ion tracks which can be acquired from the Irradiated FNTDs using WF imaging an additional interesting background signal was detected. When the exposure time was set to 10 seconds (maximum setting for the sCMOS) a unique signal appeared in all slices of the 21 z-stack layers. Fig.6 depicts the background signal found in the FNTD which may, due to the sheer number of snake like tracks be secondary electrons which transform colour centres within the FNTD. This layer is visibly above background and shows a distinct short snake-like pattern throughout the FNTD. The detection and analysis of these signals was however not further followed in this work.

## Discussion and Conclusion

LSM is currently the standard method for FNTD track readout when working with FNTDs. The ability to detect light from highly defined z-planes along with the excellent signal to noise ratio and high level of tunability situated confocal microscopy as an ideal tool for FNTD readout. However, when using the FNTD as the physical compartment in the biosensor *Cell-FIT-HD* to monitor cells for days after irradiation using WF live cell imaging the confocal system adds additional time constraints and more sources of error to the readout process, thus limiting the ability of *Cell-FIT-HD* to be a high throughput system when working with cells. The main downside of the current readout system is that two microscopes are required, one for live cell imaging (WF), the second for the FNTD readout (LSM) and therefore the FNTD needs to be transferred to a new microscope after biological imaging. The registration of track images and cell images captured on the two separate systems introduces a spatial error with regard to the ion track to cell reconstruction. As the images are captured separately on two different occasions and on two different microscopes the field reconstruction is time consuming and involves manually identifying microscopy fields by the pattern of inherent colour centre defects in the FNTD [16]. The increased time required and complexity of a dual microscope setup yields only an increase in z-axis resolution and better secondary particle detection during FNTD readout. The use of two microscopes (WF and LSM) introduces an additional error in relation to the captured FOV, which requires correction, which in turn further diminishes the use of the confocal system for FNTD readout [16].

Although the WF readout requires a single long exposure times, of up to 5s, for each image the WF system is still faster as the whole field is captured at once and no point scanning takes place. WF FNTD imaging has however been a challenge in the past as obtaining images with sufficient ion track signal intensity requires a well calibrated microscope and flatfield corrected images in addition to a sensitive modern sCMOS sensor with high quantum efficiency and a sufficiently strong illumination source.

In this paper it has been shown for the first time that is possible using a WF microscope to images the colour centres (tracks) in an FNTD. This enables a new possibility to read out the *Cell-FIT-HD* biosensor by a single WF microscope. The larger sensor enables a larger FOV which is 2.97x larger compared to the LSM images, when taken into account the WF method is therefore ∼19x faster than LSM for primary carbon ion readout. The resulting WF images from the sCMOS detector have a sufficiently high signal to background ratio which enables analysis with the existing FNTD packages developed for analysis on LSM and with far less microscope optimization steps required (scan frequency, dwell time, pinhole settings, etc.) which in turn enables a faster and more reproducible readout. The overlap of the tracks captured by imaging the same area of the FNTD using WF and LSM (Fig.4A) shows that both systems are capable of near identical outputs. The quantification of the detected track number per imaging type shows only a 1.6 % difference in primary tracks detected (Fig.4D).

Importantly for this work it needs to be stated that setting up two microscopes with different optical components, sensors (Avalanche photo diode vs sCMOS) and overall methods of illumination (point scan vs widefield) to produce the exact same image is nearly impossible. The resulting difference in captured tracks requires some level of subjective optimization of microscope settings and it depends on a user defined threshold for track detection. This value is not the same value between LSM and WF (Fig due to differences in the image datasets. Any number of hardware settings on each microscope, the definition of the minimum fluorescence intensity of the primary particle peak and the exact overlap of the FOV for each image captured per microscope add additional sources of error. Given all of these factors we believe that a value of 1.6% difference in track detection between LSM and WF is sufficient for the experiments to be performed given the large parameter space.

When comparing the expected number of primary carbon ions and their LET simulated by FLUKA (for a 333 x 333 µm area) with the detected number of particles and their energy distribution the results indicate a calculated dose difference of 6.8% between FLUKA and FNTD output for a single field using WF.

One of the possible sources for the dose mismatch could be the user defined threshold set in the FNTD analysis software during ion track analysis. It is currently not possible to remove all secondary tracks while preserving all primary tracks which leads to an altered number of tracks and ions detected which in turn will lead to a change in dose.

In addition to the decreased acquisition time when using WF there is evidence that it is now possible to detect a secondary background signal using the WF system (Fig.6), however the source of these signals still needs to be verified and quantified before it can be uniquely identified as being from any one source.

**Figure 6.**
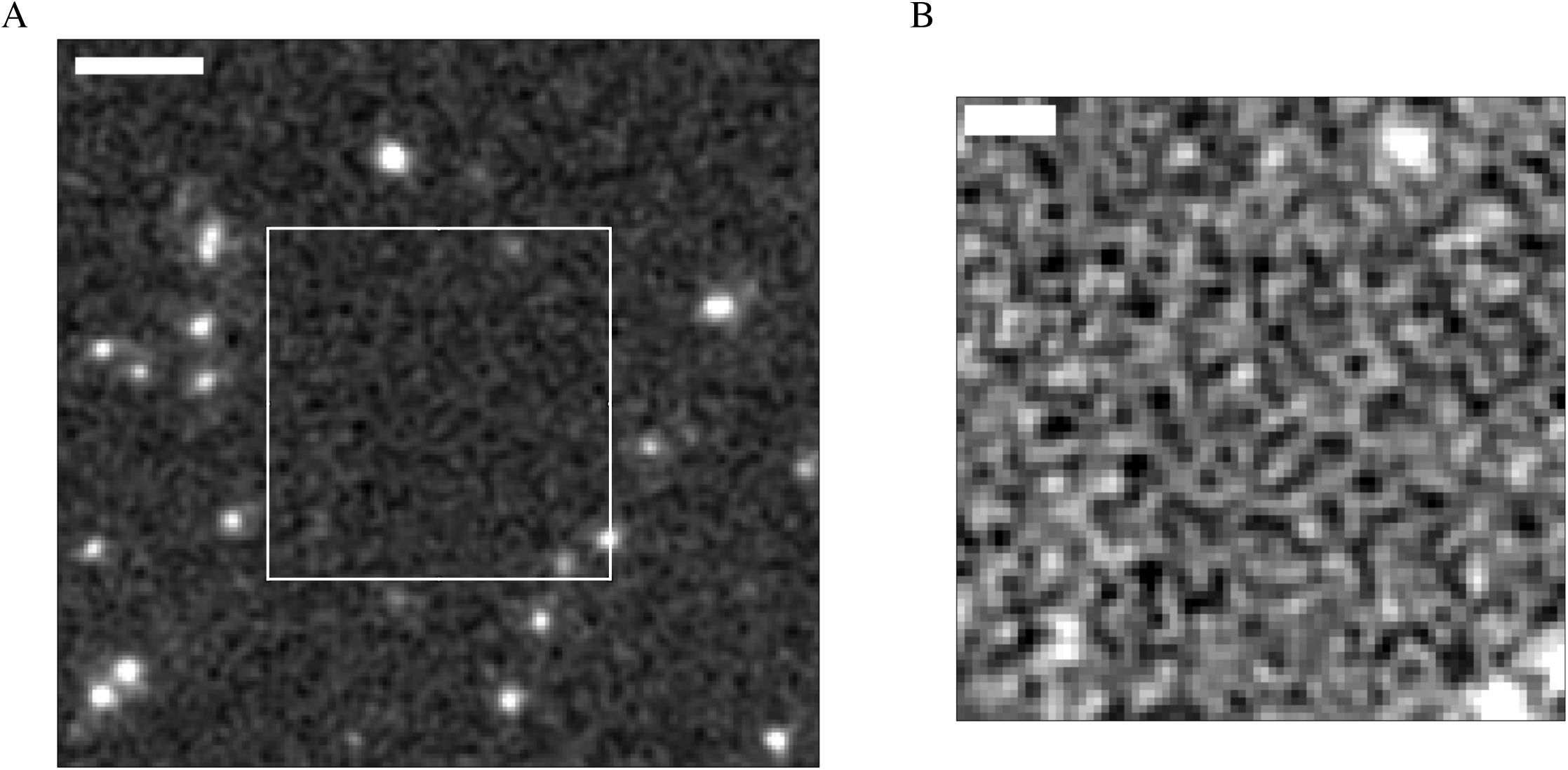
A representative depiction of the undefined background signal detected during imaging by WF microscopy. The two micrographs show the background signals detected while imaging the FNTD with the WF microscope. Left is an overview with a chosen area with a scale bar of 5µm on the right there is the selected segment with increased contrast and a scale bar of 2µm. The detected “snake” like pattern is above background and is present on all areas and z-layers captured.

In conclusion, the improvement in readout speed and the use of a WF microscope will dramatically simplify the experimental workflow of applications such as *Cell-FIT-HD*. Enabling Track analysis and consequent live cell imaging of cells for days without phototoxicity on a single imaging platform. The WF imaging protocol will enable imaging of thousands to tens of thousands of cells with their corresponding ion tracks where previously only about one hundred were possible. The images from the WF are compatible with the current analysis method for primary ion track reconstruction while requiring up to a factor ∼19 less time to acquire the initial images. The WF readout method will replace the LSM readout in *Cell-FIT-HD*. However, the WF method introduced here does not aim to replace LSM FNTD imaging in principal as a method as WF has a reduced z-resolution and cannot reach the same level of image quality that an in-depth LSM readout can. However, major advantage of WF imaging is the simple setup with comparatively few settings needed to be optimized and the dramatic decrease in time required to capture the image stacks for ion track analysis. WF FNTD imaging is an ideal solution for specific experiments, such as *Cell-FIT-HD*, where factors such as time, and the availability of two microscopes limit the microscopic analysis.

## Notes

### Competing Interest Statement

The authors have declared no competing interest.

### Summary of Updates

This revision takes into account comments from reviewers from the journal PMB. New experiments were performed and analysed to include LSM and WF measurements in the paper. Small changes to form and text were implemented. The outcome of the work however remains the same

## References

1. Akselrod, M. and J. Kouwenberg, Fluorescent nuclear track detectors – Review of past, present and future of the technology. Radiation Measurements, 2018. 117: p. 35–51.

2. Ohsawa, D., et al., Analysis of SPICE microbeam size using fluorescent nuclear track detector (FNTD). Nuclear Instruments and Methods in Physics Research Section B: Beam Interactions with Materials and Atoms, 2019. 453: p. 9–14.

3. Harrison, J., et al., CHARACTERIZATION OF FLUORESCENT NUCLEAR TRACK DETECTORS AS CRITICALITY DOSIMETERS II. Radiation Protection Dosimetry, 2017. 180(1-4): p. 201–205.

4. Dosanjh, M., The changing landscape of cancer therapy, in CERN Courier. 2018, CERN.

5. Niklas, M., et al., Subcellular Spatial Correlation of Particle Traversal and Biological Response in Clinical Ion Beams. International journal of radiation oncology, biology, physics, 2013. 87.

6. Akselrod, M.S. and G.J. Sykora, Fluorescent nuclear track detector technology – A new way to do passive solid state dosimetry. Radiation Measurements, 2011. 46(12): p. 1671–1679.

7. Osinga, J.M., et al., High-accuracy fluence determination in ion beams using fluorescent nuclear track detectors. Radiation Measurements, 2013. 56: p. 294–298.

8. Niklas, M., et al., Subcellular Spatial Correlation of Particle Traversal and Biological Response in Clinical Ion Beams. International Journal of Radiation Oncology • Biology • Physics, 2013. 87(5): p. 1141–1147.

9. Rahmanian, S., et al., Application of fluorescent nuclear track detectors for cellular dosimetry. Physics in Medicine and Biology, 2017. 62(7): p. 2719–2740.

10. Kodaira, S., et al., Co-visualization of DNA damage and ion traversals in live mammalian cells using a fluorescent nuclear track detector. Journal of radiation research, 2015. 56(2): p. 360–365.

11. McFadden, C.H., et al., Isolation of time-dependent DNA damage induced by energetic carbon ions and their fragments using fluorescent nuclear track detectors. Medical Physics, 2020. 47(1): p. 272–281.

12. Niklas M, S.J., Liew H, Zimmermann F, Dzyubachyk O,Walsh D.W.M, Rahmanian S, Holland-Letz T, Runz A, Greilich S, Debus J, Abdollahi A, The biomedical sensor Cell-Fit-HD4D, reveals individual tumor cell fate in response to microscopic ion deposition. BioRx, 2020.

13. Greilich, S., et al., Fluorescent nuclear track detectors as a tool for ion-beam therapy research. Radiation Measurements, 2013. 56: p. 267–272.

14. Dokic, I., et al., Correlation of Particle Traversals with Clonogenic Survival Using Cell-Fluorescent Ion Track Hybrid Detector. Frontiers in Oncology, 2015. 5(275).

15. Dokic, I., et al., Next generation multi-scale biophysical characterization of high precision cancer particle radiotherapy using clinical proton, helium-, carbon- and oxygen ion beams. Oncotarget, 2016. 7.

16. Niklas, M., et al., Registration procedure for spatial correlation of physical energy deposition of particle irradiation and cellular response utilizing cell-fluorescent ion track hybrid detectors. Physics in medicine and biology, 2016. 61: p. N441–N460.

17. Kouwenberg, J.J.M., et al., A 3D feature point tracking method for ion radiation. Physics in Medicine and Biology, 2016. 61(11): p. 4088–4104.

18. Team, R., RStudio: Integrated Development for R. RStudio, Inc., 2015.

19. Bauer, J., et al., Integration and evaluation of automated Monte Carlo simulations in the clinical practice of scanned proton and carbon ion beam therapy. Physics in Medicine and Biology, 2014. 59(16): p. 4635–4659.

